# A configurable model of the synaptic proteome reveals the molecular mechanisms of disease co-morbidity

**DOI:** 10.1101/2020.10.27.356899

**Authors:** Oksana Sorokina, Colin Mclean, Mike DR Croning, Katharina F Heil, Emilia Wysocka, Xin He, David Sterratt, Seth GN Grant, T Ian Simpson, J Douglas Armstrong

## Abstract

Synapses contain highly complex proteomes which control synaptic transmission, cognition and behaviour. Genes encoding synaptic proteins are associated with neuronal disorders many of which show clinical co-morbidity. Our hypothesis is that there is mechanistic overlap that is emergent from the network properties of the molecular complex. To test this requires a detailed and comprehensive molecular network model.

We integrated 57 published synaptic proteomic datasets obtained between 2000 and 2019 that describe over 7000 proteins. The complexity of the postsynaptic proteome is reaching an asymptote with a core set of ~3000 proteins, with less data on the presynaptic terminal, where each new study reveals new components in its landscape. To complete the network, we added direct protein-protein interaction data and functional metadata including disease association.

The resulting amalgamated molecular interaction network model is embedded into a SQLite database. The database is highly flexible allowing the widest range of queries to derive custom network models based on meta-data including species, disease association, synaptic compartment, brain region, and method of extraction.

This network model enables us to perform in-depth analyses that dissect molecular pathways of multiple diseases revealing shared and unique protein components. We can clearly identify common and unique molecular profiles for co-morbid neurological disorders such as Schizophrenia and Bipolar Disorder and even disease comorbidities which span biological systems such as the intersection of Alzheimer’s Disease with Hypertension.

## Introduction

At neuron-neuron synapses, the proteomes of presynaptic and postsynaptic compartments conduct neurotransmitter release and processing via complex and highly dynamic molecular networks. These signal transduction and plasticity processes underpin normal and abnormal information processing in the brain. The synaptic proteome is physically divided into pre-and postsynaptic subcellular compartments, where the postsynaptic proteome, often referred to as postsynaptic density or PSD, is believed to play a major role in processing and interpretation of the chemical messengers used in synaptic transmission between two neurons.

The PSD has been studied using high throughput proteomic methods for nearly 20 years. Initial smaller datasets (of order tens to hundreds) focused on interactomes associated with specific target proteins (NMDA receptors, AMPA receptors, PSD95, and other MAGUK proteins) [1–6]. Significant increases in the identified protein numbers (order of thousands) were achieved around 2010, when more advanced technologies were applied to whole post synaptic fractions (for example, [7–12]). To address potential noise in the sampling methods, a “consensus” PSD was proposed in 2004 [1] containing 466 proteins that had been reported in multiple studies to date and therefore were proposed as a higher confidence subset.

The pre-synaptic proteome mediates neurotransmitter release through the active zone, which organises the synaptic vesicle cycle through participation in vesicle loading, docking and priming, and calcium-triggering of fusion [13]. The history of the presynaptic proteome spans over 15 years, with the first descriptions in 2004, typically based on synaptic vesicle extraction and purification [14–18].

Proteins usually occur within supramolecular complexes and therefore proteomic datasets are often represented as static undirected protein-protein interaction (PPI) networks. Typically, the network vertices represent biomolecules, such as genes or proteins found in the studies. Edges represent the structural or protein interactions connecting them. Using such a network perspective we can exploit the many statistical measures and fundamental properties from the networks topology to gain insight and make predictions about the underlying data [19]. Vertex centrality measures such as degree and betweenness reveal influential signalling proteins within the network. The ‘scale-free’ property [20] found in many biological networks can be used to identify hub genes, which often encode disease related genes [19]. Clustering algorithms attempt to identify densely connected communities (or modules) of vertices within the network [21, 22]. Synaptic gene-disease or gene-functional association data can then be annotated onto these communities and tests made for the functional/disease enrichment at the level of the cluster. This can be used to predict gene-sets associated with known synaptic diseases, which are useful elucidating molecular mechanisms[22]. Using this network approach, we can then test where these gene-sets overlap across different disorders.

## Results

### 1. Collating the synaptic proteome

We systematically curated synaptic proteomic datasets from the literature, to produce a comprehensive index of the proteins (and their genes) reported at the synapse. To find proteomic studies we searched PubMed using the key words: “synaptic”, “postsynaptic’, “presynaptic”, “synaptosome”. Preference was given to the studies focussing on mammalian brains in healthy/normal experimental conditions. The many disease specific studies identified where not included. A total of 57 papers describing a landscape of 7814 synaptic genes (Table 1) were annotated with the following metadata: PUBMED ID, species, method of extraction, number of identified proteins and brain region. Each study’s respective protein list was extracted and mapped to stable identifiers (MGI, Entrez and Uniprot) for the predicted set of orthologues for three species (human, mouse, rat). Additional functional information (e.g. GO function, disease association) was overlaid as metadata onto the vertices. Supplementary Table 1 contains a detailed list of the papers found and associated metadata.

**Table 1:** Collection of 57 synaptic proteome studies at a glance. Postsynaptic studies are shown in blue, presynaptic in green and synaptosomal in yellow.

|  | PMID | GeneN | Brain region | Specie |
| --- | --- | --- | --- | --- |
| Husi_2000 | 10862698 | 77 | forebrain | mouse |
| Walikonis_2000 | 108118142 | 29 | forebrain | rat |
| Peng_2004 | 15020595 | 237 | forebrain | rat |
| Satoh_2002 | 11895482 | 45 | forebrain | mouse |
| Youshimura_2004 | 14720225 | 436 | forebrain | rat |
| Farr_2004 | 15447677 | 73 | whole brain | rat |
| Jordan_2004 | 15169875 | 393 | whole brain | mouse and rat |
| Li_2004 | 14532281 | 139 | forebrain | rat |
| Trinidad_2005 | 15748150 | 236 | whole brain | mouse |
| Cheng_2006 | 16507876 | 289 | forebrain and cerebellum | rat |
| Collins_2006 | 16635246 | 620 | forebrain | mouse |
| Dosemeci_2006 | 16332460 | 114 | hippocampus | rat |
| Dosemeci_2007 | 17623647 | 276 | cerebral cortex | rat |
| Trinidad_2008 | 18056256 | 2158 | cortex, midbrain, cerebellum, and hippocampus | mouse |
| Selimi_2009 | 19402746 | 63 | cerebellum | mouse |
| Fernandez_2009 | 19455133 | 301 | forebrain | mouse |
| Bayes_2001 | 21170055 | 1443 | cortex | human |
| Bayes_2012 | 23071613 | 1552 | cortex | mouse |
| Schwenk_2013 | 22632720 | 34 | whole brain | mouse |
| Distler_2014 | 25211037 | 3558 | Hippocampus | mouse |
| Bayes_2014 | 25429717 | 1134 | Frontal cortex | human |
| Frank_2015 | 2711747 | 145 | forebrain | mouse |
| Uezu_2016 | 27609886 | 928 | Cortex and hippocampus | mouse |
| Focking_2016 | 27898073 | 2021 | Whole brain | human |
| Li_2016 | 27507650 | 1602 | Hippocampus CA1 | mouse |
| Fernandez_2017 | 29045836 | 107 | forebrain | mouse |
| Roy_2017 | 29203896 | 1213 | frontal, parietal, temporal and occipital lobes of the neocortex | human |
| Li_2017 | 28671696 | 993 | Hippocampus CA1 | mouse |
| Roy_2018 | 30071621 | 1071 | frontal, medial and caudal cortex, right caudate putamen, right hippocampus, whole hypothalamus, and cerebellum (right half) | mouse |
| Wilson_2019 | 30986977 | 2134 | cortex | mouse |
| Coughenour_2004 | 15378707 | 36 | forebrain | rat |
| Blondeau_2004 | 15007177 | 209 | Whole brain | rat |
| Phillips_2005 | 16047384 | 110 | cortex | rat |
| Morciano_2005 | 16269012 | 153 | Whole brain | rat |
| Burre_2006 | 17080482 | 165 | Whole brain | rat |
| Takamori_2006 | 17110340 | 410 | Cerebral cortex | rat |
| Khanna_2007 | 17562281 | 104 | Whole brain | rat |
| Morciano_2009 | 19187093 | 369 | Whole brain | rat |
| Abul-Husn_2009 | 19562802 | 138 | Hippocampus and striatum | mouse |
| Abul-Husn_2011 | 22043286 | 145 | striatum | rat |
| Gorini_2010 | 20114047 | 57 | cortex | mouse |
| Gronborg_2010 | 20053882 | 618 | Cerebral cortex | rat |
| Boyken_2013 | 23622064 | 414 | Cerebral cortex | rat |
| Wilhelm_2014 | 24876496 | 1169 | Cortex and cerebellum | rat |
| Brinkmalm_2014 | 24973420 | 68 | Hippocampus | mouse |
| Weingarten_2014 | 24534009 | 482 | Whole brain | mouse |
| Kokotos_2018 | 30301801 | 983 | cerebellum | rat |
| Filiou_2010 | 20309889 | 2980 | Whole brain | mouse |
| Dahlhaus_2011 | 21398567 | 673 | Visual cortex | mouse |
| Cohen_2013 | 23658807 | 2668 | cortex | rat |
| Biesemann_2014 | 24413018 | 163 | forebrain | mouse |
| Chang_2015 | 25958317 | 2077 | cortex | human |
| Liu_2014 | 25352669 | 1388 | Hippocampus and prefrontal cortex | mouse |
| Distler_2014 | 25211037 | 4417 | hippocampus | mouse |
| Kohansal-Nodehi_2016 | 27115346 | 4961 | cerebellum | rat |
| Gonzalez-Lozano | 27748445 | 1560 | cortex | mouse |
| Alfieri_2017 | 28713243 | 351 | telencephalon | mouse |
| Heo_2018 | 29610302 | 2272 | Cortex and hippocampus | mouse |

#### 1.1 Post synaptic compartment

We identified 30 major post synaptic proteome (PSP) studies published in a period spanning 2000-2019 that contribute a total of 5568 mouse and human unique gene identifiers (see Supplementary Table 1). Most of these PSP datasets were obtained by proteomic analysis of biochemical fractions of synaptosomes obtained from homogenised brain samples [3–12, 23–28].

Further studies were added that focussed on specific protein complexes using immune, peptide, transgenic affinity purification or in vivo affinity purification approach strategies against well-known PSD proteins, e.g. mGluR5 [29], PSD-95 [2, 25, 26, 30], NMDA [1, 2, 31] and AMPA receptors complex [2, 32] or gephyrin [30].

The majority of PSP samples were collected from whole brain [3, 10, 29, 32, 33] and forebrain [1, 2, 4–6, 23, 27, 34, 35]. However, several studies focus on specific brain regions, e.g hippocampus [12, 24, 36, 37] and cerebellum [23, 25]. Several studies that considered multiple distinct brain regions simultaneously were also included [11, 38, 39].

The discovery rate of new PSD proteins was analysed across the multiple studies as shown in Figure 1A, where the number of proteins is plotted against the frequency of identification. Figure 1B shows the ratio of newly identified proteins (black) per year compared to the total number of PSP proteins identified in that year (blue). Two major jumps in the gross number of proteins identified occur in 2008, when 1249 new proteins were reported by [11] and in 2014 with 2588 new proteins added by [12]. The most frequently found proteins (i.e. detected in 22, or more, studies out of the 29) include very well-known PSD proteins, for example: DLG4 (found 28/29), CAMK2A (27/29), INA (26/29), SPTBN1, CAMK2B, DLG2, NSF, GRIN2B, GRIN1 (25/29), BIAP2, BSN (24/29) (full list in Supplementary Table 2).

**Figure 1.**
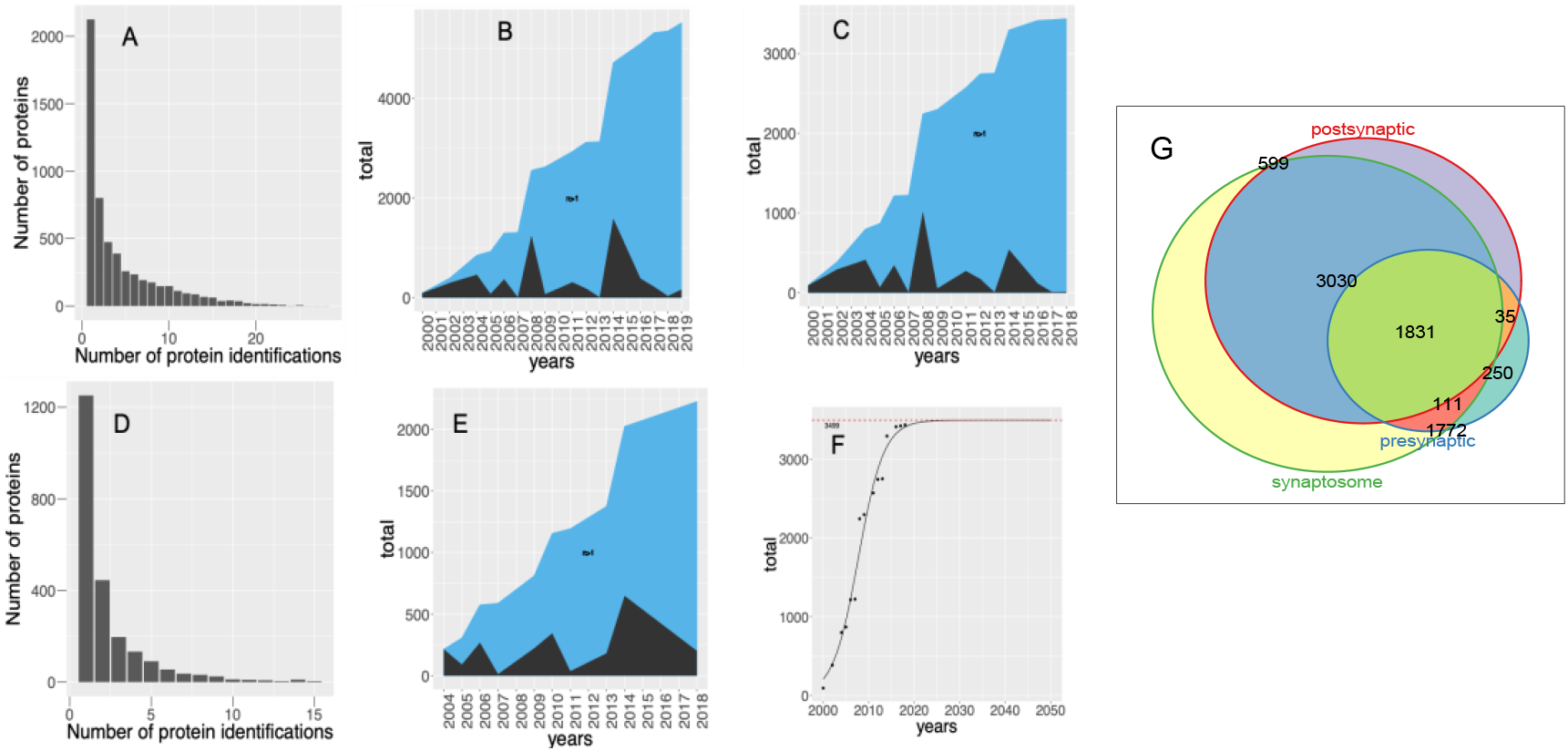
Accumulation of the unique synaptic genes by year. A. Frequency of protein identification in 29 PSP studies. B. Cumulation of the new PSP genes (black) compared to the total datasets (blue) over years. C. Accumulation of new “consensus” PSP genes (black) compared to the total datasets (blue) over years. D. Frequency of protein identification in 17 presynaptic studies. E. Accumulation of new presynaptic genes (black) compared to the total datasets (blue) over years. F. Non-linear fit to predict the total size of “consensus” PSP (3499 genes) (P = 2.36E-11). G. Overlap of three synaptic datasets: presynaptic, postsynaptic and synaptosomal.

High throughput proteomic techniques are powerful, but they are noisy, and contamination is always a concern. A large number (2091) of PSP proteins have been observed just once. While single hits may be accounted for by lack of sensitivity with low abundance molecules, it could also indicate the presence of false positive components brought in by experimental uncertainty.

The rate of growth with respect to newly discovered proteins appears to be slowing (Figure 1B) and therefore there is now an opportunity to define a more reliable subset. Following the approach described in [1], we selected genes found in two or more independent studies to designate the “consensus” PSP. This resulted in 3441 genes, which is ~7 times larger than reported by [1] and describes a subset of synaptic proteins for which have higher confidence. In this subset we observe the increment of new genes per year decreases after 2008 and drops completely after 2014 (Figure 1C). Seeing that the accumulated number of consensus PSP genes is plateauing, we performed a non-linear fit to extrapolate a predicted total number of consensus PSP genes (Figure 1F, Methods). From the fit, we predict a total number of consensus PSP genes to be 3499 (P = 2.36E-11, residual standard error: 192.7 on 12 degrees of freedom) (Figure 1F) by year 2023 which, when compared to the current number (3438) indicates that our knowledge on PSP components, based on currently available methodologies, is close to saturation.

#### 1.2 Presynaptic compartment

For the presynaptic compartment we identified 17 published studies spanning a period from 2004 to 2018, which contribute to 2315 unique human and mouse gene IDs (Supplementary Table 1). As with the PSP studies, presynaptic studies can be methodologically sorted into shotgun and target specific approaches. Shotgun approaches include the majority of studies (14/17), which span differing fractions of presynaptic compartment. This includes studies of the entire presynaptic nerve terminal and plasma membrane [16, 40, 41] to more specific studies of clathrin coated vesicles (CCV) [42], presynaptic particle fraction (PPF) [43], synaptic vesicles (SVs) [14–18, 44–47], and activity - dependent bulk endocytosis (ADBE) [48]. The three remaining studies consider target protein complexes obtained by Co-IP for N-type calcium channels (CaV2.2/CACNA1B) [49], SNAP −25 [50, 51], BKCa, dynamin-1, SNAP-25, syntaxin-1A, and VAMP-2 [51]. Presynaptic studies span a variety of anatomical regions from whole brain [15–18, 42, 49] to subregion specific, such as cortex [14, 43, 45–47, 51], hippocampus [40, 50] and striatum [40, 41].

The frequency of identification of the presynaptic proteins is shown in Figure 1D. From Figure 1E, we see two jumps in newly discovered proteins correspond to studies performed in years 2010 and 2014. Approximately half of the proteins in the presynaptic proteome (1064 genes) have been reported more than twice, which can be viewed as a rough estimate for the consensus of presynapse, however the recent trend in newly identified genes indicates that saturation has not been achieved yet (see Methods). The most frequent presynaptic genes found in the majority of the studies include AP2B1, HSPA8, GNAO1, ACTB (15/17), STX1B, ATP6V0A1, STXBP1, ATP1A3, ATP6V1E1, SYT1, GNB1, TUBA1A, VAMP2, NSF, DNM1 (14/17) with full statistics available in Supplementary Table 3.

#### 1.3 Synaptosome

Although most studies targeted pre or post synaptic regions specifically we also considered 11 studies that span the whole synaptosome and reported 6888 unique genes. These cover most but not all of the genes/proteins reported in the focussed studies (Figure 1G). In summary, 599 of specifically postsynaptic, 250 of published specifically presynaptic and 35 of their joint overlap were not detected in global synaptosome studies. Supplementary Table 4 contains the whole list of 7814 unique synaptic genes classified according their localisation (presynaptic, post synaptic, synaptosomal) based on the 57 studies considered here.

As before, most of the studies (9/11) could be classified as shotgun proteomic experiments where the entire synaptosome is analysed for its molecular components [12, 52–58]. Two studies describe the molecular structure of specific protein complexes by Co-IP targeting of kinesin (Kif5C and Kif3A) [59] and P140Cap (SRCIN1) [60]. In terms of brain region origin of the sample, the majority of the studies consider hippocampus [12, 55, 58, 59]; visual cortex [53], motor cortex [56], prefrontal cortex [59] and, cerebral cortex [61].

### 2. Reconstructing molecular interaction networks

To reconstruct protein-protein interaction (PPI) networks for the pre-and post-synaptic proteomes we used human PPI data obtained from three databases: BioGRID [62], Intact [63] and DIP [64]. These data were filtered for the highest confidence direct and physical interactions (see Methods; Table 2), for which we excluded all predicted interactions and those obtained from Co-IP experiments. The resulting PSP network contains 4,817 nodes and 27,788 edges in the Largest Connected Component (LCC). The presynaptic network is significantly smaller and comprises 1780 nodes and 6620 edges in the LCC.

**Table 2.** Summary of protein-protein interactions (PPIs) in this study. PPI numbers before and after filtering for direct and physical interactions (Methods)

| Total N of unique Human PPIs extracted from three databases | N of unique Human PPIs after filtering for direct/physical interaction | N of unique PPIs in PSP network | N of unique PPIs in PSP "consensus" network | N of unique PPIs in Presynaptic network |
| --- | --- | --- | --- | --- |
| 407,643 | 126,579 | 28,915 | 14,496 | 6,625 |

### 3. Using the Synaptic Proteome Model

We captured the original heterogeneity of the datasets by incorporating the necessary metadata so that they can be used in queries that retrieve specific subsets of interest, e.g. only human, only presynaptic, or, say, only genes obtained in Co-IP experiments from hippocampus. To make the datasets readily accessible based on custom selection criteria we embedded them in a SQLlite relational database.

In addition to the proteomic, interactomic data and metadata mentioned above, the database also includes the GO function information for three species: mouse, rat and human, disease annotation for human (based on Human Disease Ontology (HDO)[65]) and GeneToModel table, which links certain synaptic proteins to existing computational models of synaptic plasticity and synaptic signal transduction (*Sterratt in preparation*). The database structure is shown at Figure 2 and described in detail in respective Methods section.

**Figure 2.**
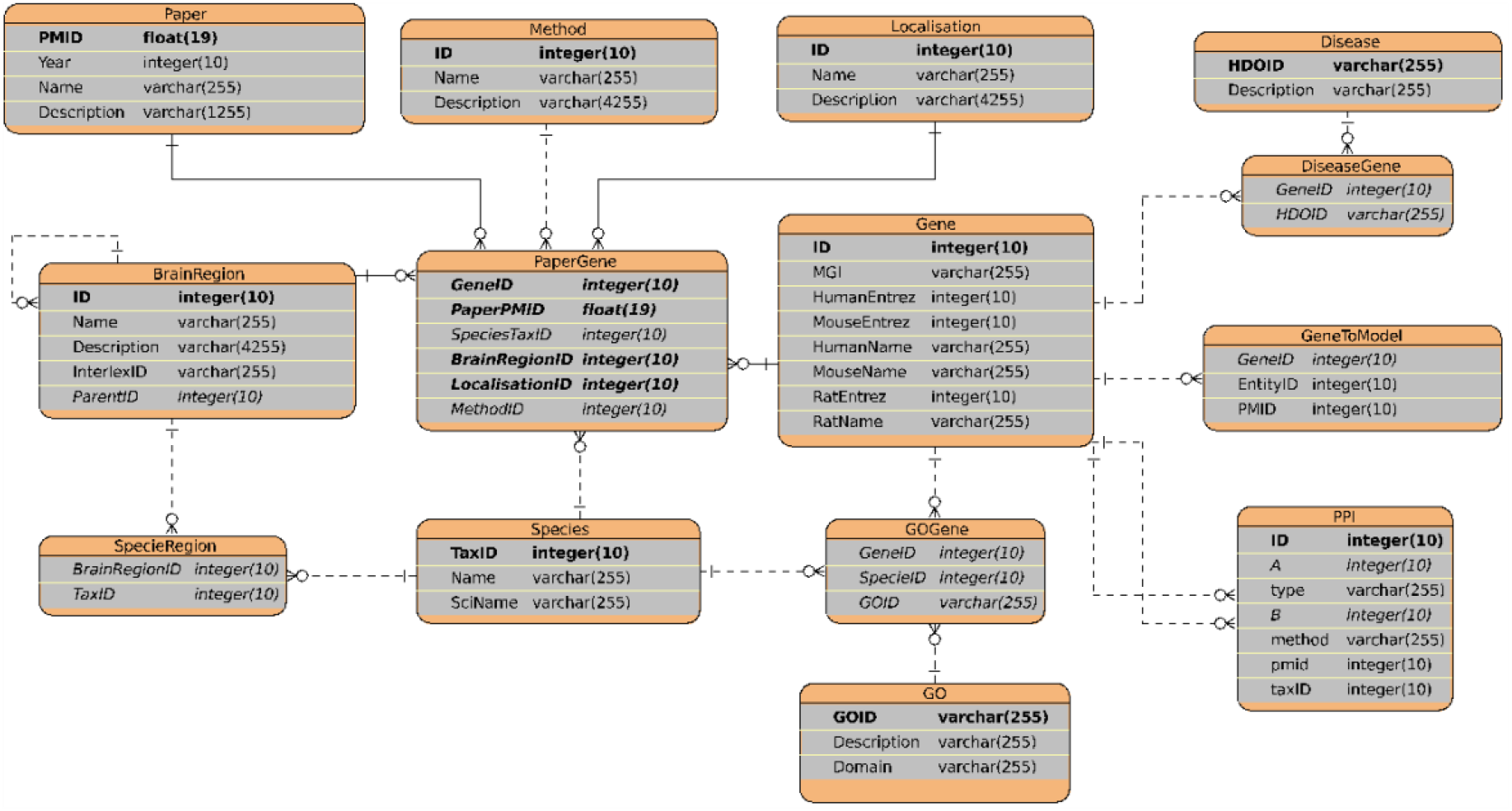
Entity Relationship diagram for the SQLite database of the synaptic proteome.

We provide the data as simple flat tables for maximum flexibility and a SQLlite implementation. For anyone who wishes to explore the data we also provide a simple walkthrough of how to install the system and run the following common uses/queries from SQLite Studio or R (Supplementary Files). Following use cases illustrate how the database can be used to obtain the information for specific protein(s) and build the PPI network for subset of proteins (1,2), and how the PPI network can be used for further analysis (3,4).

Database is available from Supplementary Data.

#### 3.1. Finding information for specific gene/s of interest

Among the common questions in the neuroscientific community are “What is known about my favourite gene/protein? Is it pre- or postsynaptic? Which brain region it was found and by whom?”. Evidence to address these types of questions can be extracted by simple database queries using protein name (gene name) or gene Entrez ID. For example, to find the information for protein “SRCIN1”, the query will look like:

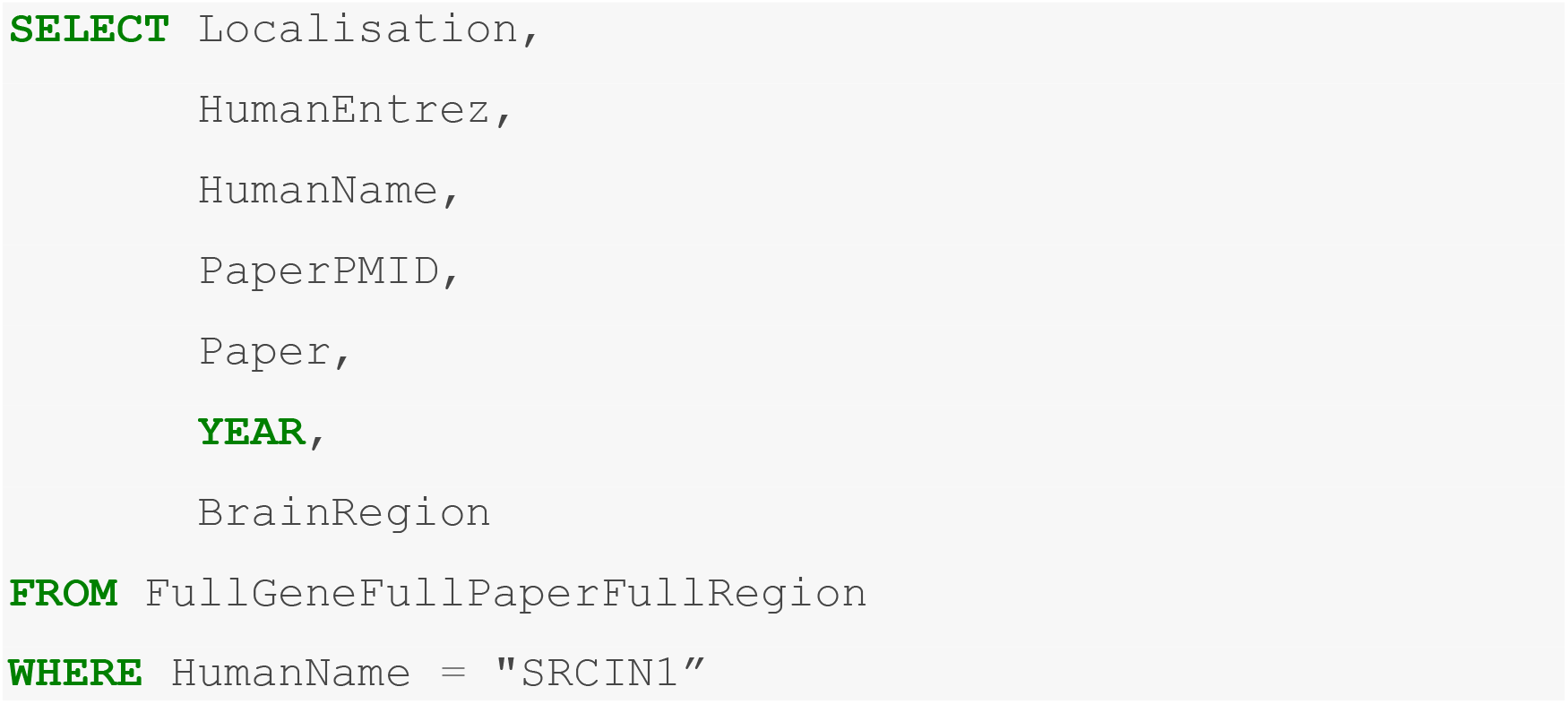

Here, FullGeneFullPaperFullRegion is a database view that combines the major information for all genes in more convenient spreadsheet style representation.

The database returns the results in a form of table (Table 3 contains top 17 rows), showing that SRCIN1 was found in 25 postsynaptic, 7 presynaptic and 12 synaptosomal studies, along with brain regions where it was reported.

**Table 3.** A snippet of available information for SRCIN1

| Compartment | EntrezID | GeneName | PMID | AUTHOR_YEAR | Brain region |
| --- | --- | --- | --- | --- | --- |
| Postsynaptic | 80725 | SRCIN1 | 15447677 | FARR_2004 | Brain |
| Postsynaptic | 80725 | SRCIN1 | 15020595 | PENG_2002 | Forebrain |
| Postsynaptic | 80725 | SRCIN1 | 18056256 | TRINIDAD_2008 | Midbrain |
| Postsynaptic | 80725 | SRCIN1 | 18056256 | TRINIDAD_2008 | Cerebellum |
| Postsynaptic | 80725 | SRCIN1 | 30071621 | ROY_2018 | Hypothalamus |
| Postsynaptic | 80725 | SRCIN1 | 16332460 | DOSEMESI_2006 | Hippocampus |
| Postsynaptic | 80725 | SRCIN1 | 30071621 | ROY_2018 | Striatum |
| Postsynaptic | 80725 | SRCIN1 | 17623647 | DOSEMESI_2007 | Cerebral cortex |
| Postsynaptic | 80725 | SRCIN1 | 29203896 | ROY_2017 | Frontal lobe |
| Presynaptic | 80725 | SRCIN1 | 24876496 | WILHELM_2014 | Cerebellum |
| Presynaptic | 80725 | SRCIN1 | 17110340 | TAKAMORI_2006 | Cerebral cortex |
| Synaptosome | 80725 | SRCIN1 | 27115346 | KOHANSAL_NODEHI_2016 | Cerebellum |
| Synaptosome | 80725 | SRCIN1 | 28713243 | ALFIERI_2017 | Telencephalon |
| Synaptosome | 80725 | SRCIN1 | 25211037 | DISTLER_2014 | Hippocampus |
| Synaptosome | 80725 | SRCIN1 | 27115346 | KOHANSAL_NODEHI_2016 | Cerebral cortex |
| Synaptosome | 80725 | SRCIN1 | 25352669 | LIU_2014 | Prefrontal cortex |
| Synaptosome | 80725 | SRCIN1 | 25958317 | CHANG_2015 | Motor cortex |
| Synaptosome | 80725 | SRCIN1 | 21398567 | DAHIHAUS_2011 | Visual cortex |

#### 3.2. Building a custom PPI network from a protein list

We can easily extract a customised subset of the PPI network that includes specific studies, brain regions or compartments. The example query for a presynaptic protein list and its respective PPI network is as follows:

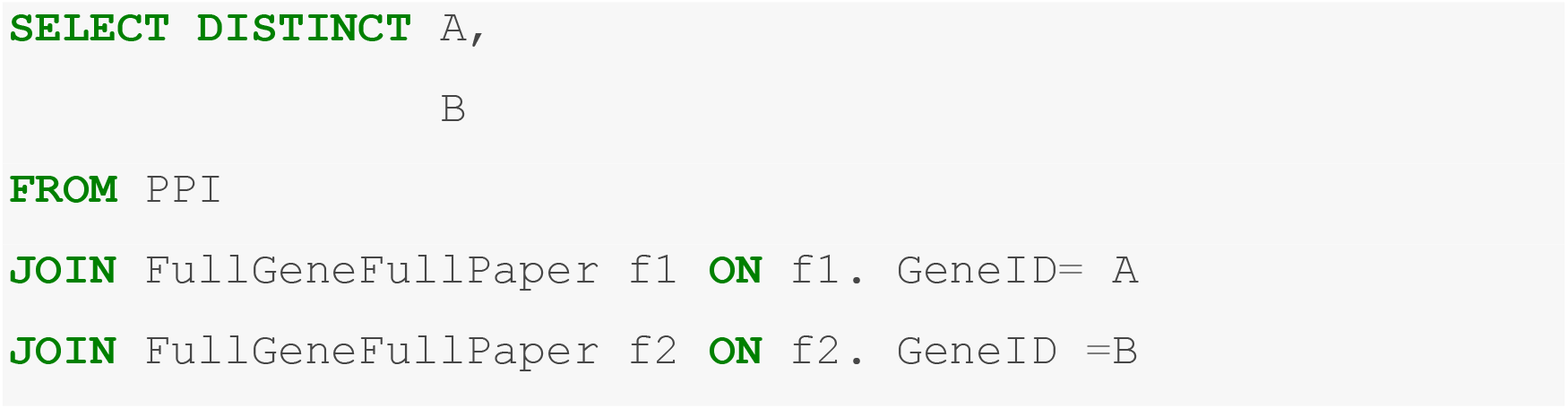
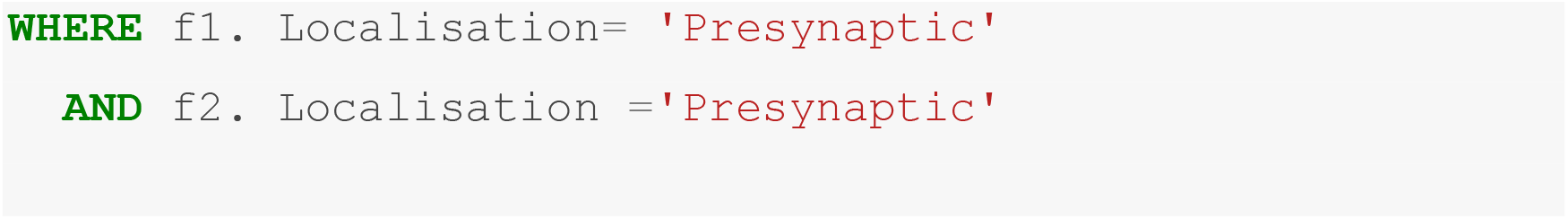

This will result in a list of protein interaction pairs of type “Human_Entrez_ID_A - - Human_EntrezID_B”, which can be visualised in R, using package *Igraph* (https://igraph.org/redirect.html), loaded to Cytoscape via *RCy3* package [66] or saved as csv file and analysed with Cytoscape (https://cytoscape.org/) or Gephi (https://gephi.org/).

#### 3.3. Measuring how network topology correlates with functional/disease importance: Bridgeness

The relative importance of specific proteins to propagate information (i.e. signals) through a network can be estimated from the position of the protein within the network’s architecture and its substructures. The number of proteins interacting neighbours (degree) is considered as important property as it may reflect protein importance for signal propagation in the network. This can be extended to include local interacting partners, a measure known as Semi-local centrality [67].

Clustering is commonly used to identify substructures/communities within networks. Concrete clustering methods usually assign each protein to the single most likely community despite many of them actually being involved in multiple communities. To match this we calculate the probability of each protein being involved in every cluster in the network. The more communities a specific protein “bridges” (has a probability belonging to) the more likely it is involved in communicating between these communities across the network [68] (Methods). We hypothesise these ‘Bridging’ proteins will also correlate with functional/disease importance at the biological level.

Using the Spectral clustering algorithm [71], we estimated the Bridgeness for the PSP network and identified a total of 3884 (3884/4817 ~ 80%) proteins with Bridgeness value more than 0.1.

We then plotted Bridgeness against Semi-local centrality values (see Methods) to categorise the proteins by their ability to influence the network globally and locally (Figure 3, Supplementary Table 5). Region 1 contains 1126 proteins (see Region 1 in Figure 3, Supplementary table 5), which, between them affect many (i.e. greater than half) communities in the network yet each is relatively localised in the network (low local-centrality values) – we call these “bridging” or “influential”. Of those 1126 −28 are considered as core PSD95/DLG4 interactors [34] (28/77 ~36%) within the postsynaptic proteome including: DLG4, GRIN2A, GRIN2B, CAMK2A, CALM1, NSF. Interestingly, we found NMDA-receptor subunits proteins GRIN1 and AMPA-receptor subunit proteins (GRIA2,3,4) clustered together as secondary Bridging proteins (see Region 3 in Figure 3), indicating their influence on a small number of selected functional clusters. There are a few scaffold proteins which bind to the AMPA-receptor subunits which are Bridging Proteins, e.g. DLG1, PICK1, GRIP1.

**Figure 3.**
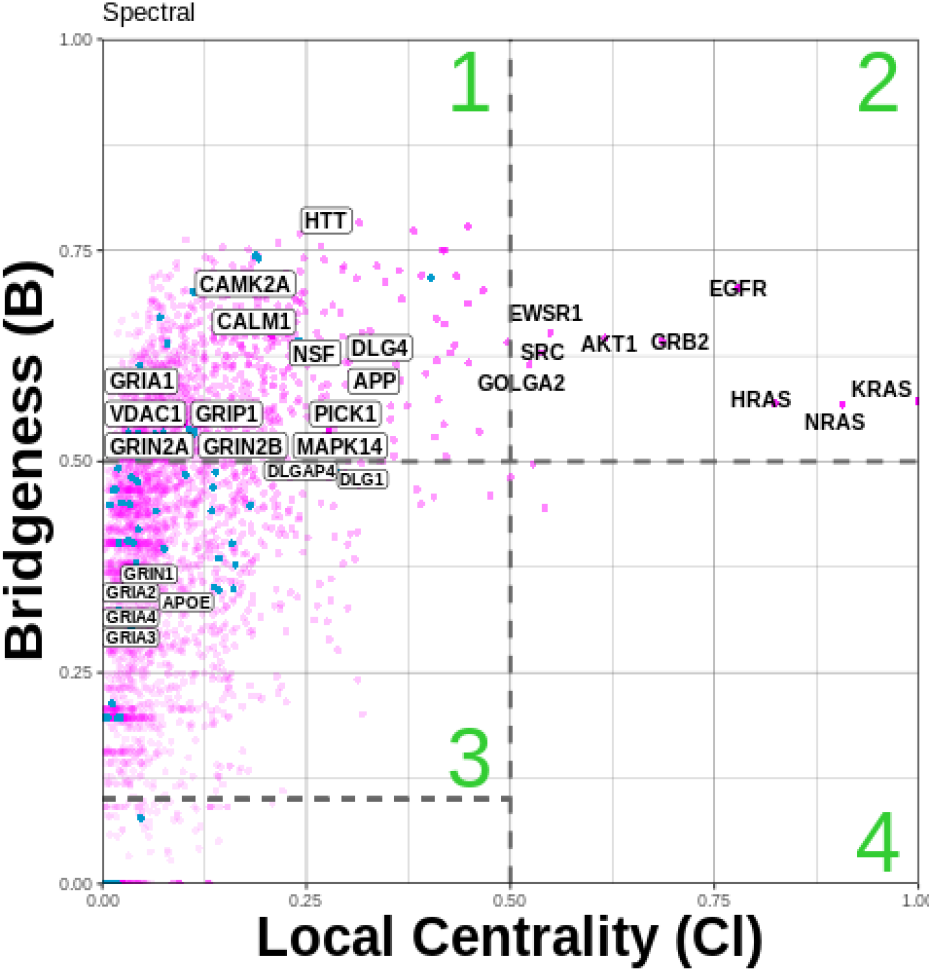
Bridging proteins estimated using the Spectral clustering algorithm. Bridgeness is plotted against semi-local centrality (Methods), allowing categorisation of the proteins to three important groups: Region 1, proteins that are likely to have a ‘global’ rather than ‘local’ influence in the network (also been called bottle-neck bridges [69], connector or kinless hubs [70] lie in the range (DLG4, GRIN2B, CAMK2A, etc). Region 2, proteins likely to have an influential ‘globally’ and ‘locally’ in the network (EGFR, HRAS, NRAS, etc). Region 3, more likely to be centred within the community they're found in, but also focused on communicating with a few other specific communities (GRIN1, GRIA2-4), “secondary bridging proteins”. Region 4, proteins whose impact is mostly ‘locally’, primarily within one or two communities (local hubs [68]).

Among 1126 Bridging proteins, 512 (512/1126 ~45%) were found associated with at least one synaptopathies given in our set (Methods), 248 (214/1126 ~ 19%) with multiple diseases including: APP (AD&Epi&ASD&PD&HTN&MS&FTD), VDAC1 (AD&PD&MS), and MAPK14 (AD&SCH&HD&HTN&MS). Indeed, using the Spectral algorithm we found significant overlaps between the Bridging proteins (see Region 1 in Figure 3) and disease association; AD (P=7.74E-10,***), HTN (P=1.16E-07***), HD (P=8.01E-7***), PD (P=7.9E-5***). Moreover, when we considered synaptic function (GO annotation), we found a significant overlap for AD with innate immune response-activating signal transduction (GO:0002758, AD=4.3×10-11, HTN=9.7×10-10), stress-activated MAPK cascade (GO:0051403, AD=3.310-12, HTN=4.3×x10-11) and calcium-mediated signaling (GO:0019722, AD=1.1×10-3).

#### 3.4. Extracting and comparing shared disease pathways

Given gene-disease annotation (GDA) data and the PPI network topology, we can start to dissect how these different neurological diseases might coalesce at the synapse. The methodology described in [72] tests the extent to which two diseases might share a common molecular mechanism. Briefly, within the network we calculate z-scores from the observed mean shortest distance between disease GDA’s, relative to 10000 random permutation of each diseases GDA’s, to quantify the disease pairs overlap (Methods). From the set of p-values, calculated from these z-scores, we then correct for false positives by calculating the q-values (Supplementary Table 6).

Figure 4 shows the comparison of the disease pair overlaps obtained for common neurological and neurodevelopmental diseases and disorders in pre- and post-synaptic networks. It can be seen that the overlap for common neuro-psychiatric/developmental disorders was observed in both postsynaptic and presynaptic PPI network models (post/pre), however q-values are smaller for presynaptic network, probably due to its size: BP-ASD (P=8.79E-14/1.44E-03), BP-SCH (P=2.65E-25/9.55E-11), ASD/SCH (P=8.74E-07/9.82E-03). Similarly, overlap was observed for common neurodegenerative diseases/conditions AD and PD (P=1.74E-5/1.92E-05).

**Figure 4.**
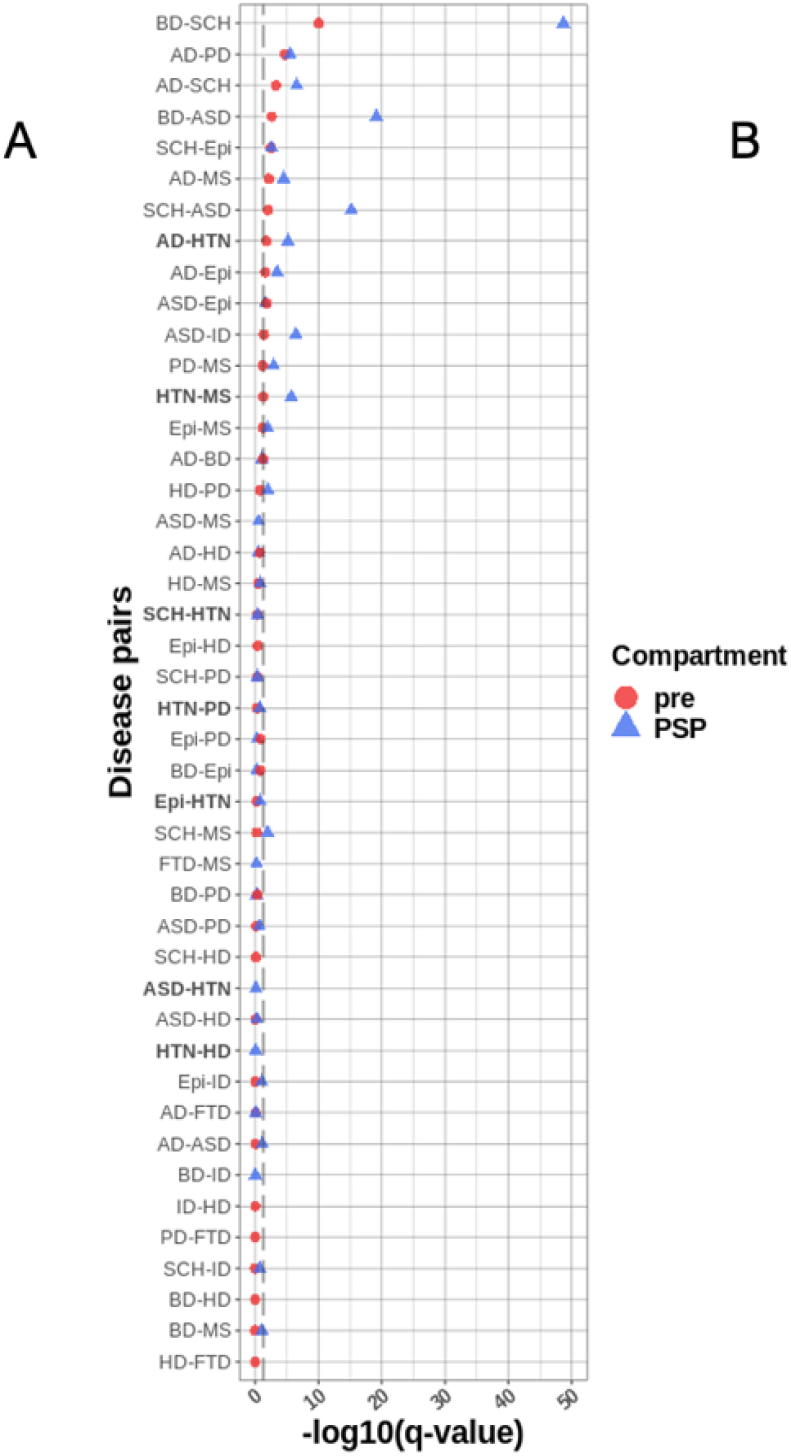
Disease-disease relationship for presynaptic and postsynaptic interactome. Where significance q-values < 0.05 is delineated by the dashed line. Schizophrenia (SCH), Autistic Spectrum Disorder (ASD), Autistic Disorder (AUT), Bipolar Disorder (BD), Intellectual Disability (ID), Alzheimer disease (AD), Epilepsy Syndrome (Epi), Parkinson’s Disease (PD), Frontotemporal Dementia (FTD), Huntington’s Disease (HD) and Multiple Sclerosis (MS) are considered.

Hypertension (HTN) is considered one of the leading vascular risk factors associated with neurological diseases/disorders including AD, PD and FTD [73–75], and BD and SCH [76, 77]. With the PSP PPI network model, we found a strong overlap for HTN with AD (P=8.6E-4, and with MS (P=8.79E-5), but not with the other neurodegenerative diseases: FTD-HTN, PD-HTN, Epi-HTN and HD-HTN which all have q-values > 0.05 (Supplementary Table 6). When considering the set of common neuro-psychiatric/ developmental disorders with HTN, we found all four overlap with HTN, i.e., but that there was no evidence these overlaps were significant when considering a full randomised model: HTN-SCH (P=0.9), HTN-BD (P=0.9), HTN-ID (P=0.32) and HTN-ASD (P=1).

The more a disease pair overlaps, the greater the evidence the corresponding disease pair share the same genes, and therefore participate in similar pathways. Since we find significant overlaps between AD-HTN and AD-PD, but not PD-HTN, it indicates a potential shared mechanistic pathway between AD and HTN, which is different to the pathways shared between AD and PD; Supplementary Figure 3A and 3C illustrates this difference by plotting the z-score calculated against the distribution of for AD-HTN and PD-HTN after 10000 randomised models using the PSP network.

## Discussion

By systematic interrogation of the published literature we identified 57 studies that describe a molecular landscape of the synaptic proteome comprising 7617 unique mouse and human genes. Of this, the majority of the papers (29/57) are for the postsynaptic compartment, 17 are presynaptic and 11 consider the whole synaptosome. We combined all the datasets based on stable identifiers, which resulted in 5568 postsynaptic genes (3441 found in two or more studies forming an updated “consensus” PSP), 2315 presynaptic genes, and 6888 reported for synaptosome. We retrieved the PPIs for the list of genes, which allows us to build a comprehensive PPI map for the whole synapse or its subcellular compartments.

We embedded the datasets in a SQLite database to enable more flexible queries based on the species, brain region, method of identification or other metadata and allow the extraction of bespoke protein network models for subsequent analyses.

This dataset is the largest and the most complete up to date and is freely available with lightweight tools to allow anyone to extract relevant subsets. By mirroring the methods used it would be straightforward for any user to add in their own datasets for comparison.

The case studies described provide useful examples of the analyses that can be readily extracted from these data. In addition, to information for specific proteins, we can use the model to analyse the entire protein-protein interaction network to query topology-function relationships that provides insights for possible disease mechanism. For example, since the Bridging proteins are spread out over the network, the correlated disease-disease pairs we find are not centred in a single cluster but shared across several communities. In turn, the co-occurrence of enrichment of specific synaptic functions with disease in the discovered communities may indicate that molecular complexes underlying the specific synaptic functions are also involved in disease mechanisms. When comparing across diseases we find evidence for shared molecular mechanisms that span common neurological disorders, such as Bipolar disease and Autism Spectrum Disorder, Bipolar Disease and Schizophrenia. Moreover we see evidence for molecular mechanisms that span more diverse disease pairs such as the existence of common molecular pathways linked to both Alzheimer’s disease and Hypertension.

## Methods

### 1. Data cleaning and ID mapping

Synaptic studies for presynaptic, postsynaptic and whole synaptosome compartment were identified by manual curation of PubMed, starting from year 2000. Protein/gene lists from the published synaptic proteome studies were combined into the total list of synaptic components. Identifiers from each study were mapped to stable IDs including: Entrez Human and Mouse, Uniprot and MGI IDs. The resulting ‘master list’ contains only unique gene entries (rows) with corresponding studies (columns) where they have been identified. For example, if a protein was discovered in several studies all those respective columns will contain 1. A summary of all studies is presented in Supplementary Table 1. It contains the information for literature ID and some useful metadata such as species (mouse, human, rat), protein counts, brain region where the sample was taken from along with experimental methods used for protein identification and quantification.

### 2. Prediction of total size of pre- and postsynaptic proteome

Only proteins that were found more than one time were taken into account to make the most confident “consensus” dataset for pre-and post-synaptic proteomes. We fitted the accumulation of new proteins against the year they were first time identified in R, using linear (y ~ x), and non-linear (y ~SSlogis (x, Asym, xmid, scal)) models. The goodness of fit was compared by Akaike’s Information Criteria) AIC function [78], where lower indicates a more parsimonious model.

For post synaptic proteome the non-linear model is shown at Figure 1F and predicted maximum size of “consensus” PSP proteome is 3499, achieved by roughly 2023. AIC coefficient is 213.8955 for linear fit and 205.0504 for non-linear fit, which means the latter is more parsimonious.

For presynaptic proteome the non-linear fit is shown at Supplementary Figure 1, predicting 1309 proteins in total reached by 2035 year.

However, by AIC criteria, the liner model for presynaptic “consensus” proteome is better than non-linear (103.2001 and 107.2766, respectively), which likely means that presynaptic proteome is not in its “saturation” phase yet.

### 3. Protein-protein interaction data

Protein-Protein Interaction (PPI) dataset was assembled from three publicly available databases: DIP [62]. IntAct [61] and BioGRID [60]. The interactions were retrieved in PSI-MITAB standard protein interaction format [74, 75]. The PSI-MITAB format enforces unified column contents and use of controlled vocabularies. These columns report different types of identifiers of an interactor pair, the PubMed identifier of publication where the interaction was reported, taxonomies of interactors, an interaction detection method, and an interaction type. The last two identifiers are defined with unique MI-terms specified in the Molecular Interactions Controlled Vocabulary (MI) ontology. The set of MI-terms representing each of the categories was obtained by parsing the MI ontology file (source: https://www.ebi.ac.uk/ols/ontologies/mi).

To limit the analysis to high-confidence direct physical interactions, those annotated by “association” term (MI:0914) and its offspring were preserved. This step excluded interactions of types: colocalization, functional association, genetic interaction and predicted interaction. Although MI ontology offers “direct interaction” term in the category of interaction type, according the the iMEX curation rules (https://github.com/IMEx-Consortium/IMEx-site/raw/gh-pages/static/files/imex_curation_rules.pdf) two hybrid assays are categorised as non-direct interactions. Therefore, this classification was not used to preserve interactions reported by two-hybrid assays. We further removed indirect interactions, often linked to methods likely to generate false positive hits (spoke-expansion). Interactions originating from experiments involving co-complexes (e.g. pull-down, affinity technology) were excluded from the analysis by filtering out a selected subset of terms denoting these methods. Since the intAct interaction table indicates spoke-expanded interactions, these were also removed from the interaction set.

Integration of interaction tables required distinct parsing procedures per database, including translation into common interactor identifiers. DIP and IntAct databases use UniProtKB accessions (UniProtKB ACs) as their primary identifiers, e.g. P49418. On the other hand, BioGRID uses numeric Entrez Gene IDs, e.g. 1134. The unification of identifiers was achieved by combining mapping tables collected from NCBI and UniProtKB ftp servers (file addresses: ftp://ftp.uniprot.org/pub/databases/uniprot/current_release/knowledgebase/idmapping/by_organism/RAT_10116_idmapping_selected.tab.gz ftp://ftp.uniprot.org/pub/databases/uniprot/current_release/knowledgebase/idmapping/by_organism/HUMAN_9606_idmapping_selected.tab.gz ftp://ftp.uniprot.org/pub/databases/uniprot/current_release/knowledgebase/idmapping/by_organism/MOUSE_10090_idmapping_selected.tab.gz ftp://ftp.ncbi.nlm.nih.gov/gene/DATA/gene2accession.gz). Entrez Gene ID was chosen as a unifying identifier.

Interactors that were annotated with a different taxon than Mouse, Rat or Human were removed. Non-human Entrez Gene IDs were mapped to Human identifiers with gene orthology mapping tables available on NCBI ftp server (file address: https://ftp.ncbi.nlm.nih.gov/gene/DATA/gene_orthologs.gz).

After application of the above criteria, 407 443 interactions were obtained. Additionally, after manual curation, 200 interactions from PDB database were added directly to the PPI list with Method “MI:0114” – X-ray crystallography, and Type “MI:0407”-Direct interaction, which resulted in total of 407643 interactions.

All files where retrieved on the same date, 2020-06-18, parsed and integrated with a custom code available at https://gitlab.com/Wysocka/ppis_v02.

### 4. Estimation of Bridgeness and semi-local centrality measures

To assess the topological importance of the proteins in PSP network we estimated two independent measures for each protein.

1) semi-local centrality Cl(v), which takes into consideration both a vertex’s degree, its nearest, and next to nearest neighbours:

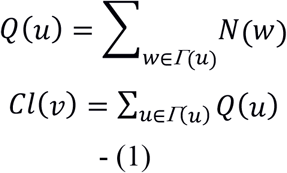

where *Γ(u)* is the set of nearest neighbours of *u* and *N(w)* is the number of nearest and next to nearest neighbours of the vertex *w*. We performed unity-based or feature scaling: X – Xmin/(Xmax-Xmin) to normalise the semi-local centrality to lie in the range [0,1].

Semi-local centrality differs from degree centrality therefore, in making use of more information, allowing us to also measure a vertices’ spread’ of information locally through the network.

2) Bridgeness *B(v)* of vertex *v* to measure the influence of a gene due to the clustering [68] can be estimated as:

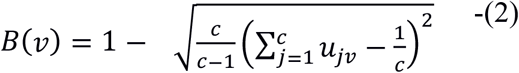

Where *u*_*v*_ is the community membership vector for vertex *v*, which is the probability of vertex v belonging to a given community *u*_*v*_ = [*u*_1*v*_, *u*_2*v*_, ..., *u*_1*cv*_], where ∑_*j*_*u*_*jv*_ = 1 and c is the number of communities detected by the algorithm.

The combination of these measures allows categorisation of the influence each has on the overall network structure (Figure 3):

### 5. Disease Network Localisation

We compared the gene-disease association for the pre and post-synaptic PPI networks using the combined OMIM/GeneRIF/Ensembl variation data for common set of synaptic diseases (or synaptopathies) [79]: Schizophrenia (SCH, DOID:5419), Autistic Spectrum Disorder (ASD, DOID:0060041), Autistic Disorder (AUT, DOID:12849), Bipolar Disorder (BD, DOID:3312), Intellectual Disability (ID, DOID:1059), Alzheimer disease (AD, DOID:10652), Epilepsy Syndrome (Epi, DOID:1826), Parkingson’s Disease (PD, DOID:14330), Frontotemporal Dementia (FTD, DOID:9255), Huntington's Disease (HD, DOID:12858), Multiple Sclerosis (MS, DOID:2377) and Hypertension (HTN, DOID:10763).

We investigated the overlap and separation of each disease-disease pair by measuring the mean shortest distance for each disease, using the shortest distance between each GDA to its next nearest GDA neighbour [72]. The overlap, or separation, of each disease-disease pair in the pre-post-synaptic PPI networks, could then be quantified using:

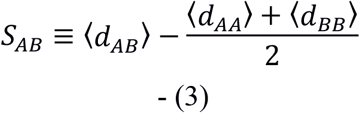

Where 〈*d*_*AA*_〉and 〈*d*_*BB*_〉 quantify the mean shortest network distance between genes associated with disease A (or B), and 〈*d*_*AB*_〉 the mean shortest distance between diseases. *S*_*AB*_ is bound by the diameter of the network, i.e., *d*_*max*_ ≤ *S*_*AB*_ ≤ *d*_*max*_ where *d*_*max*_ is 8, 7, 8 for the presynaptic, PSP and PSP consensus PPI networks respectively. The magnitude of *S*_*AB*_ depends on the number of GDSs associated with each disease. Large positive values imply two well separated diseases, while large negative values indicate large (number of GDAs) diseases with a big overlap, often implying one disease is the variant or precursor to the other. Each disease-disease network separation pair (*S*_*AB*_) was compared against a full randomised model: drawing the same number of GDAs (from the set of all network genes) for each disease at random, before computing its separation. For each disease-disease pair, we performed 10,000 iterations of the full randomised model using the ECDF distributed computing facility.

The difference between the observed and randomised disease pair separations, was quantified using the z-score:

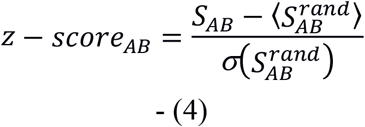

Where 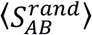 and 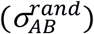are the mean and standard deviation obtained from the 10,000 iterations. Each disease-disease pair separation using the full randomised model, i.e., 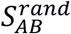, was found to follow a normal distribution. We therefore assessed the significance of each disease-disease pair’s separation, from P-values estimated from its z-score calculated in (4):

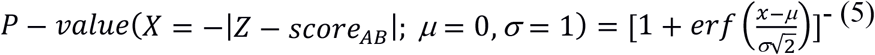

Where we take the negative of the absolute value of each disease-disease pairs z-score calculated in (4) and make use of R’s *pnorm* function available in the ‘stats’ package (R version 3.4.2).

The confidence in each disease-disease pairs P-value was tested for by calculating its q-value [80], and from the Bonferroni correction at the 0.05 (*), 0.01 (**) and 0.001 (***) significant levels.

### 5. SQL database

The database contains the following main tables:

- Gene: list of genes including IDs (MGI, Entrez Human and Mouse) and gene names (Human and Mouse).
- Specie: Tax ID (Human and Mouse)
- Paper: list of papers with PMID ID, name (in format “FirstAuthor_year”), year of publication
- Location: postsynaptic, presynaptic, synaptosome
- Method: shotgun or IP
- Brain region: list of regions where the samples originate, with hierarchical region structure
- PPI: human protein-protein interactions combined from BioGRID, Intact and DIP databases, contain information on methods, interaction type (PSI-MI nomenclature) and PMID info for each of the interactions.
- PaperGene: table links gene to respective papers and the metadata above
- GO: BP, CC and MF GO annotation for Human, Mouse and Rat species
- GOGene: gene to GO association list
- Disease: List of diseases from the HDO for Human
- DiseaseGene: genes to disease association list
- GeneToModel: genes with found association with published model of synaptic plasticity

The database is created with SQLite v_3.31.1_ RDBMS in SQLite Studio v3.2.1.

## Supporting information

Synaptic proteome database

## Acknowledgements

OS, CM, JDA, DS, were funded by European Union Horizon 2020 Specific Grant Agreement No. 72027 (Human Brain Project SGA2). OS and JDA were funded by European Union Horizon 2020 Specific Grant Agreement No. 945539 (Human Brain Project SGA3). The work of S.G.N.G. was supported by the UK Medical Research Council (G0802238, Defining the Human Synapse Proteome) and Simons Initiative for the Developing Brain. The Simons Initiative for the Developing Brain (SIDB) at the University of Edinburgh is supported by the Simons Foundation Autism Research Initiative (529085)

## Author contribution

OS, CM, MDC selected and curated the datasets; EW, KFH, XH, TIS designed and performed the PPI extraction and filtering; CM and OS performed the network analysis, wrote the analysis scripts and performed the statistical testing; OS constructed the SQLite database; OS, CM, JDA, DS, TIS, SGNG designed and directed the study, wrote and edited the manuscript.

## Supplementary Figures and Tables

Figure 1. Non-linear model for accumulation of presynaptic genes over years (year of first identification is considered), predicted number of 1309 genes, which should be reached by 2035.

Figure 2. The hierarchical structure of brain regions (human) existing in the database, according to 57 selected studies.

Figure 3. Linear mapping of Z-scores with ecd function used for assigning of gene compartment localisation.

Supplementary Table 1. List of 57 synaptic publications combined for the database with information on year, PUBMID, gene count, method, brain region and short description.

Supplementary Table 2. List of 5568 post synaptic genes collected from 29 publications listed as columns. 1 or 0 in each row corresponds to “found” or “not found” for each specific gene in each specific publication.

Supplementary Table 3. List of 2315 presynaptic genes collected from 17 publications listed as columns. 1 or 0 in each row corresponds to “found” or “not found” for each specific gene in each specific publication.

Supplementary Table 4. Full list of 7814 unique synaptic genes classified according their localisation (presynaptic, post synaptic, synaptosomal) based on 57 studies considered here. Localisation gets “1” if at least one publication has reported this gene for this compartment.

Supplementary Table 5. Estimation of Bridgeness values for PSP network protein.

Supplementary Table 6. Estimation of disease pairs overlap.

